# Promise and Limits of Hierarchical Dynamical RNNs for Individualized Resting-State fMRI

**DOI:** 10.64898/2026.03.20.713153

**Authors:** Carlotta B.C. Barkhau, Keyvan Mahjoory, Manuel Brenner, Elias Weber, Ramona Leenings, Clemens Pellengahr, Nils R. Winter, Maximilian Konowski, Tabea Straeten, Susanne Meinert, Elisabeth J. Leehr, Kira Flinkenflügel, Tiana Borgers, Dominik Grotegerd, Hannah Meinert, Julia Hubbert, Christoph Jurischka, Judith Krieger, Wiebke Ringels, Frederike Stein, Florian Thomas-Odenthal, Paula Usemann, Lea Teutenberg, Igor Nenadic, Benjamin Straube, Nina Alexander, Andreas Jansen, Hamidreza Jamalabadi, Tilo Kircher, Markus Junghöfer, Udo Dannlowski, Tim Hahn

**Affiliations:** Institute for Translational Psychiatry, University of Münster, Germany; Institute for Biomagnetism and Biosignalanalysis, University of Münster, Germany; Ernst Strüngmann Institute for Neuroscience, Frankfurt, Germany; Dept. of Theoretical Neuroscience, Central Institute of Mental Health (CIMH); Charité – Universitätsmedizin Berlin, Department of Psychiatry and Neuroscience, Berlin, Germany; Institute for Translational Neuroscience, University of Münster, Germany; Department of Clinical Psychology and Psychotherapy Georg-Elias-Müller-Institute of Psychology, Georg-August-University of Göttingen, Germany; Department of Psychiatry and Psychotherapy, University of Marburg, Germany; Center for Mind, Brain, and Behavior (CMBB), University of Marburg, Germany; Faculty of Medicine, University of British Columbia, Canada; Bielefeld University, Medical School and University Medical Center OWL, Protestant Hospital of the Bethel Foundation, Department of Psychiatry, Bielefeld, Germany

**Author notes:** **Corresponding author:** Carlotta B.C. Barkhau.

## Abstract

Modeling individual brain dynamics from resting-state fMRI (rs-fMRI) remains challenging due to substantial inter-subject variability, noise, and limited data length per subject. Here, we systematically evaluate whether hierarchical shallow piecewise-linear recurrent neural networks (shPLRNNs), recently introduced as interpretable dynamical system reconstruction models, can generate individualized rs-fMRI time series while preserving subject-specific functional connectivity structure.

We applied the framework to 1,344 rs-fMRI samples from healthy participants of the Marburg-Münster Affective Disorders Cohort Study (MACS). Simulated rs-fMRI data reproduced substantial empirical FC structure, with comparable reconstruction accuracy on the validation and held-out test sets. Generalization to unseen individuals was heterogeneous and strongly depended on how typical a subject’s connectivity pattern was relative to the training cohort, with template similarity explaining 37% of variance in reconstruction accuracy.

Learned subject-specific parameters exhibited significant test-retest stability and higher within-subject than between-subject similarity on longitudinal data from two different timepoints, supporting their interpretation as individualized dynamical markers. Associations between individual parameters and demographic or cognitive variables were statistically significant but modest in effect size, and predictive performance remained below that obtained using empirical rs-fMRI features directly. Empirical FC was used as a reference for static subject information rather than as a target to be outperformed. Together, these results suggest that hierarchical shPLRNNs can extract meaningful and partially stable individual-specific dynamical structure from rs-fMRI data. The findings delineate key trade-offs between model expressivity, generalization and subject specificity, and point to directions for future methodological refinement in individualized brain modeling.

## 1. Introduction

Understanding individual differences in brain function is a central aim in neuroscience, with important implications for personalized medicine and psychiatric research. Resting-state functional magnetic resonance imaging (rs-fMRI) has become a widely used tool for probing intrinsic brain activity, providing non-invasive access to functional organization and connectivity patterns. A central challenge, however, is that individual brain dynamics must be inferred from relatively short, indirect, and motion-sensitive BOLD time series (1). This makes it difficult to distinguish stable subject-specific structure from measurement variability and population-level regularities (2,3).

A range of methodological approaches has been developed to improve individual-level inference from rs-fMRI. Precision fMRI approaches (4) and individual-specific parcellation techniques (5–7) aim to better characterize subject-specific functional topographies, while functional connectome fingerprinting and predictive modeling have demonstrated that empirical functional connectivity (FC) contains substantial individual-specific information (8,9). At the same time, whole-brain modeling (10–12) and machine learning frameworks tailored to noisy, high-dimensional data (13–15), provide (mechanistic) accounts of how large-scale neural dynamics may give rise to observed FC patterns. These approaches have substantially advanced individualized neuroimaging, but they also emphasize different aspects of the problem (16,17): empirical FC is a powerful static summary of subject-specific covariance structure, mechanistic models offer biological interpretability, and predictive machine learning approaches are often optimized for downstream classification or regression rather than for reconstructing generative brain dynamics.

Temporal deep learning models, including recurrent neural networks (RNNs), offer a complementary route for modeling rs-fMRI as a dynamical process. Recent RNN-based and graph-based approaches have been used to forecast or simulate fMRI activity by learning temporal dependencies in multivariate brain signals (18–20). However, many such approaches focus primarily on prediction accuracy or sequence forecasting. For individualized neuroscience, an additional question is whether a model can learn a shared dynamical structure across subjects while preserving stable, subject-specific differences in a compact and interpretable form.

From this perspective, hierarchical RNNs are particularly relevant. Rather than treating each subject independently, hierarchical architectures leverage shared information across a cohort while representing individual differences through low-dimensional subject-specific parameters (20,21). This structure is well suited to rs-fMRI, where per-subject time series are limited but group-level data are abundant. Importantly, hierarchical RNNs can be understood not only as flexible sequence models, but also as tools for dynamical system reconstruction: they aim to infer an underlying generative process whose long-term behavior gives rise to observed time series (19,20). In this setting, successful modeling is not defined solely by pointwise prediction accuracy, but by the ability to generate trajectories that preserve emergent properties of the empirical system, such as subject-specific FC structure and temporal dependencies.

The shallow piecewise-linear recurrent neural network (shPLRNN) provides a compact and interpretable framework for such dynamical system reconstruction (DSR). Recent work has shown that hierarchical shPLRNNs can recover shared and individual-specific dynamics from time series data across subjects and experimental conditions, while yielding low-dimensional latent representations that can be related to underlying control parameters (20,22). Whether this framework can be successfully applied to high-dimensional empirical rs-fMRI data remains an open question. In particular, it is unclear how well hierarchical shPLRNNs can preserve individual FC structure, how their performance generalizes to unseen subjects, and whether the learned subject-specific parameters contain stable and phenotypically meaningful information.

Here, we systematically evaluate hierarchical shPLRNNs for individualized modeling of rs-fMRI dynamics in the Marburg-Münster Affective Disorders Cohort Study (MACS). Our goal is not to replace empirical FC as a static fingerprinting or prediction tool, but to assess what a compact generative dynamical model can recover from short rs-fMRI time series. Specifically, we ask: first, whether simulated trajectories preserve empirical FC structure across training and held-out subjects; second, whether generalization depends on how closely an individual’s FC resembles the population-level connectivity template; third, whether subject-specific model parameters are stable across repeated sessions; and fourth, whether these parameters are associated with demographic and cognitive variables such as sex, age, body mass index (BMI), intelligence quotient (IQ), and years of schooling (YoS). By explicitly evaluating both strengths and limitations, this study aims to clarify the potential of hierarchical recurrent dynamical models for individualized rs-fMRI.

## 2. Methods

### 2.1 Dataset, acquisition, and data split

Data were drawn from the Marburg-Münster Affective Disorders Cohort Study (MACS), a two-site neuroimaging cohort collected in Marburg and Münster, Germany, using identical study protocols and harmonized scanner settings (23,24).

The functional imaging protocol included an approximately 8-min resting-state fMRI sequence. Resting-state fMRI was acquired using a T2*-weighted echo-planar imaging sequence with repetition time (TR) = 2000 ms, echo time (TE) = 30 ms in Marburg and 29 ms in Münster, field of view = 210 mm, matrix size = 64 × 64, slice thickness = 3.8 mm, distance factor = 10%, flip angle = 90°, anterior-to-posterior phase encoding direction, ascending acquisition order, 33 slices, and an effective voxel size of 3.28 × 3.28 × 4.18 mm³.

The study was approved by the ethics committees of the medical faculties at the University of Marburg and the University of Münster. All participants provided written informed consent and received financial compensation. At the time of analysis, resting-state fMRI data were available from 953 healthy control subjects. For a subset of 491 subjects, a second rs-fMRI session was available approximately two years after the first measurement, resulting in 1,444 rs-fMRI samples in total.

Samples with at least 200 usable time points after preprocessing and motion censoring were retained, yielding a final dataset of 1,344 samples. All time series were truncated to a common length of 200 time points to provide fixed-length inputs for the hierarchical shPLRNN framework.

To prevent data leakage, all samples from the same subject were assigned exclusively to one data split. The final dataset was divided into a training set (n = 945), a validation set used for hyperparameter selection (n = 208), and a held-out test set used only for final evaluation (n = 191) (Figure 1).

**Figure 1:**
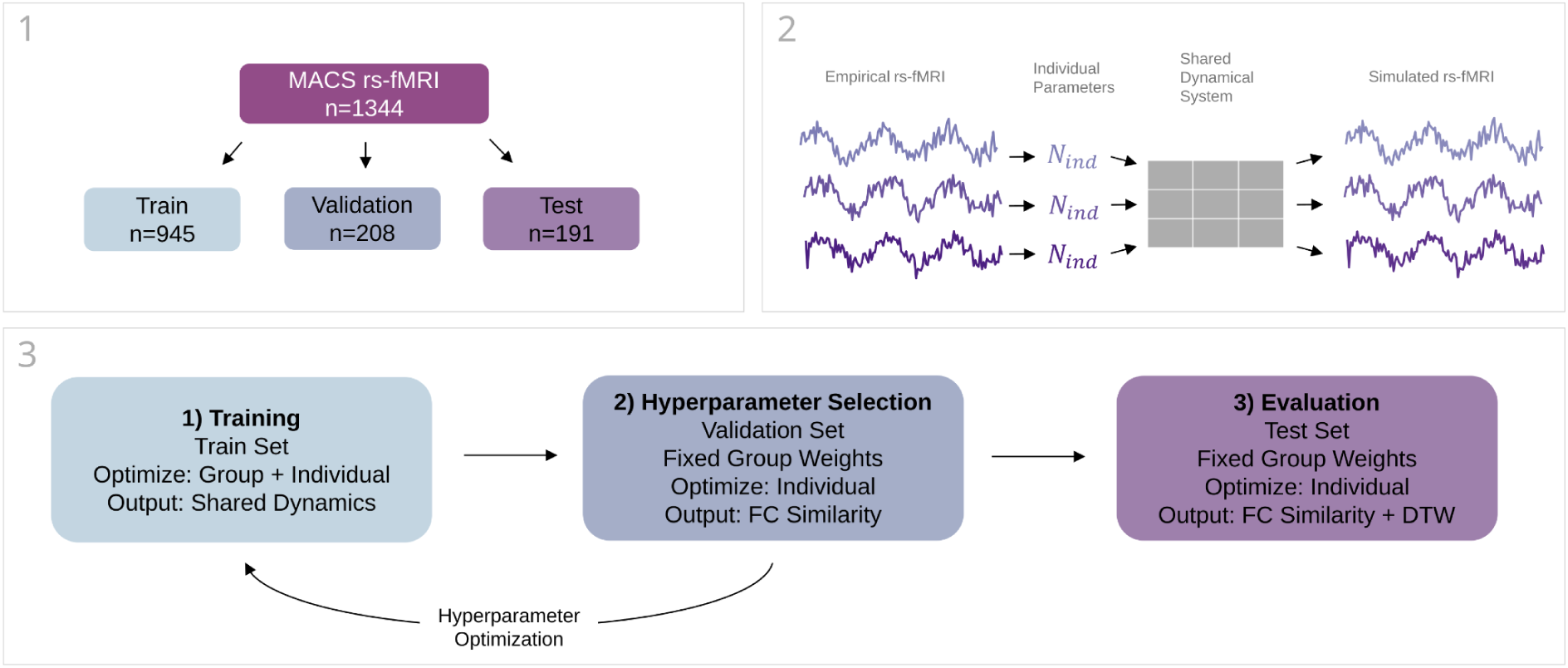
Study overview and modeling workflow. 1) Subject-wise split of the MACS resting-state fMRI dataset into training, validation and held-out test sets to prevent data leakage. 2) Conceptual illustration of the hierarchical shPLRNN architecture with shared group-level recurrent dynamics and low-dimensional subject-specific parameters. 3) Training, hyperparameter selection and evaluation procedure. Group-level and individual parameters are jointly optimized on the training set. Hyperparameters are selected on a separate validation set while keeping group-level parameters fixed. Final performance is assessed on a held-out test set after subject-specific fine-tuning.

### 2.2 rs-fMRI preprocessing and functional connectivity

Preprocessing of empirical rs-fMRI data was performed using the open-source Connectivity Analysis Toolbox (CATO) (25). The preprocessing pipeline included slice timing correction using FSL sliceTimer, motion correction using FSL MCFLIRT (26), and anatomical parcellation based on each participant’s T1-weighted image. Mean BOLD time series were extracted for cortical regions of interest defined by the Desikan-Killiany atlas, resulting in 68 cortical ROI time series per sample.

The Desikan-Killiany atlas is an anatomical cortical parcellation comprising 68 cortical regions. We selected this relatively low-dimensional atlas to obtain stable ROI-level time series for dynamical modeling of short rs-fMRI sequences and to reduce the dimensionality of the observed system relative to the available sequence length. This choice favors robustness and computational tractability but may limit sensitivity to fine-grained individual functional topography.

Motion-related metrics, including framewise displacement (FD) and DVARS, were computed for each frame following Power et al. (2). A zero-phase Butterworth band-pass filter between 0.01 and 0.1 Hz was applied to retain low-frequency fluctuations commonly analyzed in resting-state fMRI while avoiding phase distortions (27). Frames exhibiting excessive motion were censored using the following thresholds: maxFD = 0.25, maxDVARS = 1.5, minViolations = 2, backwardNeighbors = 1, and forwardNeighbors = 0. A frame was flagged only when both FD and DVARS exceeded their thresholds, and each flagged frame was removed together with its preceding frame. Censored frames were removed rather than interpolated, and the remaining frames were concatenated. Only samples with at least 200 remaining frames were retained, and the first 200 frames were used for modeling. Across the final 1,344 samples, 23.3% contained no censored frames, and the median number of removed frames was 5 (95th percentile: 19; maximum: 23). Because each censoring event removes at least two consecutive frames, splice points affect at most 1.6% of one-step transitions. The influence of censoring on reconstruction accuracy is assessed in Supplement S7.

FC matrices were computed as Pearson correlation coefficients between the mean BOLD time series of all pairs of ROIs. For all analyses involving FC similarity, matrices were vectorized by extracting the upper triangular elements excluding the diagonal.

### 2.3 Hierarchical shPLRNN model

We applied a hierarchical DSR framework based on shallow piecewise-linear recurrent neural networks (shPLRNN) to model ROI-level rs-fMRI time series across subjects (20). In contrast to conventional time-series forecasting models, DSR models aim to learn generative surrogate models of the underlying dynamical process, such that freely generated trajectories preserve relevant long-term temporal and geometrical properties of the observed system.

Each rs-fMRI sample was represented as a multivariate time series

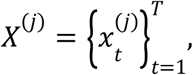

with *T* = 200 time points and 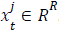, where *R* = 68 denotes the number of cortical regions of interest. The model therefore receives and generates the full multivariate ROI-level signal, rather than modeling each ROI independently.

The hierarchical framework assumes that each subject is described by a subject-specific dynamical system, while sharing a common group-level parameterization across subjects. Subject-specific differences are represented by a low-dimensional learnable feature vector 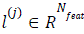. This feature vector is mapped to the full set of subject-specific DSR parameters 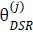 through a shared group-level mapping 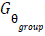:

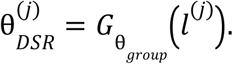

In this way, the model separates population-level structure, encoded by the shared group-level mapping, from individual variability, encoded by the low-dimensional subject-specific feature vectors.

Following Brenner et al. (20), the subject-specific dynamics were implemented using a shallow piecewise-linear recurrent neural network. In all fMRI experiments, we used the clipped shPLRNN variant, in which the nonlinear term is implemented as a difference of rectified linear activations. The latent state 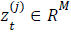 evolves according to

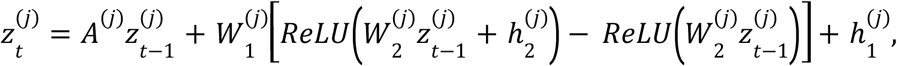

where *A*^(*j*)^ is a subject-specific diagonal matrix, 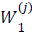 and 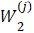 are subject-specific recurrent weight matrices, 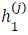 and 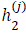 are subject-specific bias terms, and ReLU denotes the rectified linear unit. The piecewise-linear form of the model makes the learned dynamics mathematically tractable while retaining sufficient flexibility to approximate nonlinear dynamical systems.

In the present application, the latent dimensionality was set equal to the number of observed ROIs (*M* = *R* = 68), and the observation function was taken to be the identity mapping, such that generated latent states directly corresponded to simulated ROI-level rs-fMRI signals:

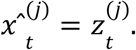

Subject-specific parameters were generated from the feature vector *l*^(*j*)^ by linear projection through shared group-level parameter matrices. Thus, the low-dimensional subject-specific feature vector determines the individual dynamical regime, while the shared matrices define how feature vectors are translated into full RNN parameters.

The resulting model can be interpreted as a compact generative representation of individual rs-fMRI dynamics. Its goal is not to forecast future BOLD activity from a fixed input horizon, but to learn subject-specific dynamical systems whose generated trajectories reproduce emergent properties of the empirical data, in particular functional connectivity structure. See Figure 1 for a schematic overview.

### 2.4 Training objective and hyperparameter selection

Model training was performed end-to-end using generalized teacher forcing (GTF), a training procedure designed to stabilize recurrent dynamical system reconstruction by linearly interpolating between model-generated and data-inferred latent states during optimization (28). In our implementation, the teacher-forcing coefficient was gradually reduced from 0.5 to 0.15 over training. GTF was used only during optimization; model evaluation was based on autonomously generated trajectories.

During initial training, shared group-level parameters and subject-specific feature vectors were optimized jointly on the training set. The model was trained on multivariate ROI-level rs-fMRI subsequences sampled from the preprocessed 200-time-point trajectories. For each training sample, the input sequence consisted of consecutive time points, and the target sequence was shifted forward by one time point. Thus, training optimized one-step-ahead sequence reconstruction under GTF, while final evaluation was based on freely generated trajectories. During initial training, subsequences of length 30 were used. During subject-specific fine-tuning, subsequences of length 100 were used.

The training objective was implemented as a Gaussian negative log-likelihood with diagonal covariance:

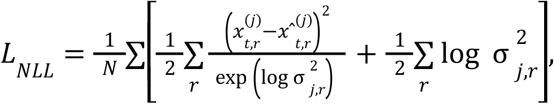

where 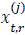 denotes the empirical signal of subject *j*, ROI *r*, and time point 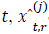 denotes the corresponding model-generated signal under GTF, and log 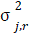 denotes the subject-specific diagonal log-covariance term. The loss was averaged across training samples.

Functional connectivity similarity was not directly optimized during training but was used only for hyperparameter selection and model evaluation.

Optimization was performed using Adam with separate optimizers for shared group-level parameters and subject-specific quantities. The learning rate was set to 1×10^−4^ for shared group-level parameters and 1×10^−3^ for subject-specific parameters. Both learning rates were decayed using an exponential scheduler with factor 0.999 per epoch. Weight decay was applied only to shared parameters and was set to 0.

Two architectural hyperparameters were optimized using Optuna (29): (i) the number of subject-specific feature parameters *N_feat_* ∈ {20, 50, 100, 500}, and (ii) the hidden size *L* of the shPLRNN, sampled between 250 and 1360. The latent dimensionality was fixed to the observation dimensionality (*M* = *R* = 68). Models were trained with a batch size of 2,000 subsequences. For each batch, subsequences were sampled from a randomly selected subset of 250 subjects, corresponding on average to eight subsequences per selected subject.

For each hyperparameter configuration, the model was first trained on the training set for 1,000 epochs. Group-level parameters were then kept fixed, and subject-specific fine-tuning was performed on the validation set for an additional 1,000 epochs, during which only subject-specific quantities were updated. The hyperparameter configuration minimizing 1 − *r_FC_*, where *r_FC_* denotes the mean Pearson correlation between empirical and simulated FC matrices after fine-tuning, was selected for all subsequent analyses.

After training or fine-tuning, simulated rs-fMRI trajectories were generated autonomously for each subject. Simulations were initialized with the first empirical ROI-level time point of the corresponding subject. For evaluation, 210 time points were generated, the first 10 time points were discarded as burn-in, and the remaining 200 time points were used to compute simulated FC matrices and temporal similarity measures.

### 2.5 Model evaluation and robustness analyses

Model performance was quantified primarily by the Pearson correlation between empirical and simulated FC matrices. For each subject, empirical and simulated FC matrices were vectorized by extracting the upper triangular elements excluding the diagonal, and Pearson correlation was computed between the resulting FC vectors. This metric assesses whether generated trajectories reproduce the subject-specific FC structure of the empirical rs-fMRI data.

As a complementary temporal similarity measure, we computed dynamic time warping (DTW) distance between empirical and simulated multivariate time series. DTW allows local stretching or compression of the time axis to align similar temporal patterns despite timing shifts (30). Prior to DTW computation, each ROI time series was z-scored separately for empirical and simulated data so that DTW reflects temporal structure rather than signal amplitude. DTW was computed along the time axis and divided by the number of time points, yielding a distance per time point (lower values indicate higher similarity). DTW was included as a secondary measure, because the primary modeling objective was the reproduction of emergent dynamical properties such as FC rather than exact pointwise temporal prediction. Note that simulated trajectories are generated autonomously and contain no splice points, whereas empirical time series may contain a small number of discontinuities due to censoring (Section 2.2). FC, as a zero-lag measure, is unaffected by frame order. DTW may be slightly affected, which is a further reason we treat it as a secondary measure, see Supplement S7 for further investigation.

To assess the contribution of population-level connectivity structure to individual reconstruction performance, we first averaged empirical FC matrices across the training set to obtain a group-mean FC template. For each held-out test subject, we then computed three Pearson correlations: (i) between individual empirical FC and the group-mean FC, serving as a static template baseline; (ii) between simulated FC and the group-mean FC, quantifying the similarity of model-generated FC to the population template; and (iii) between simulated and empirical FC of the same subject, representing individual reconstruction accuracy. To contextualize the template baseline, we additionally computed a noise-matched baseline, defined as the mean correlation with the FC of each individual training subject (i.e., the expected similarity to a single randomly chosen training scan), and compared each subject’s simulated FC with their own versus other subjects’ empirical FC.

To assess inter-individual variability in generalization performance, we further examined whether template similarity predicted reconstruction accuracy. Using linear regression, we quantified how much of the between-subject variance in reconstruction accuracy could be explained by similarity to the group-mean FC template, thereby assessing whether the model generalized better to subjects whose FC patterns were more typical of the training cohort.

To contextualize model performance, we conducted several robustness analyses. First, we trained a reduced hier-shPLRNN variant with only one subject-specific parameter per individual, limiting the model’s capacity to capture individual differences (Supplement S1). Second, we repeated model training using progressively smaller training subsets of 500, 100, and 50 subjects to assess dependence on training cohort size (Supplement S2). Third, we repeated the full training procedure five times with independent random initializations and compared the resulting simulated FCs across runs to assess run-to-run stability (Supplement S3). Fourth, we related the number of censored frames to reconstruction accuracy (Supplement S7).

### 2.6 Subject specificity and test-retest reliability

To assess whether simulated FC retained subject-specific information beyond group-level structure, we performed an FC-based subject identification analysis within the training set (9,31). For each subject, the simulated FC matrix was correlated with all empirical FC matrices in the training set. We then recorded whether the empirical FC of the matching subject achieved the highest correlation with the simulated FC (Top-1), ranked among the three highest correlations (Top-3), or ranked among the five highest correlations (Top-5). This analysis was used to quantify the extent to which generated trajectories preserve subject-discriminative FC structure.

To assess test-retest stability and subject specificity of hier-shPLRNN parameters, we used the subset of training subjects with two rs-fMRI sessions acquired approximately two years apart. For each session, subject-specific model parameters were estimated independently while keeping the group-level parameters fixed. Subjects with two available sessions in the training set were included in this analysis (n = 308).

Parameter similarity between two sessions was quantified as the Pearson correlation between the full subject-specific parameter vectors. Prior to similarity computation, parameter values were z-standardized across subjects to account for scale differences. As a reference baseline, we performed an analogous analysis on empirical FC. FC matrices were vectorized by extracting the upper triangular elements excluding the diagonal, and test-retest FC similarity was defined as the Pearson correlation between corresponding FC vectors.

For each subject, within-subject similarity was computed between the two sessions. Between-subject similarity was estimated using a matched procedure: for each within-subject comparison, one session from a different subject was randomly selected, yielding equal numbers of within– and between-subject comparisons. Statistical significance of within-subject similarity exceeding between-subject similarity was assessed using a one-sided permutation test with 20,000 permutations. A one-sided Mann-Whitney U test was used as a non-parametric robustness check. Effect sizes were quantified using Cohen’s d and Cliff’s delta.

Finally, we assessed whether model-level stability was related to data-level stability by correlating subject-wise parameter test-retest similarity with empirical FC test-retest similarity using Pearson and Spearman correlation coefficients.

### 2.7 Phenotypic association analyses and baseline representations

We next assessed whether subject-specific hier-shPLRNN parameters retained phenotypic information. We examined associations with sex, age, body mass index (BMI), years of schooling (YoS), and intelligence quotient (IQ). Analyses were restricted to the first available time point per subject to avoid data leakage across repeated measurements, and to the training set to avoid using held-out test subjects for secondary model interpretation, resulting in n = 606 subjects for the phenotypic analyses.

For univariate statistical analyses, we tested each subject-specific parameter separately. Sex was analyzed using analysis of variance (ANOVA), while age, BMI, YoS, and IQ were analyzed using ordinary least squares (OLS) regression, similar to analysis in (32). All models included age, sex, and scanning site as covariates, except when the variable served as the outcome itself. For continuous outcomes, the general regression model was

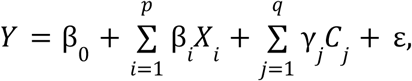

where *Y* denotes the target variable, *X_i_* denotes the *i*-th subject-specific model parameter, *C_j_* denotes the *j*-th covariate, and ε denotes the residual error.

For the ANOVA on sex, we included age and scanning site as covariates and quantified effect size using partial eta-squared 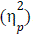. For continuous outcomes, standardized beta coefficients (β*_std_*) were used as effect sizes. Multiple comparisons across subject-specific parameters were controlled using the Benjamini-Yekutieli false discovery rate correction with q < 0.05 (33). For each target variable, the parameter showing the strongest association was selected for visualization and effect-size reporting. Bootstrap confidence intervals were computed using the bias-corrected and accelerated (BCa) bootstrap method, with group stratification where applicable.

To provide a multivariate estimate of predictive information, we additionally performed machine learning analyses using PHOTONAI (13,34,35). Pipelines included missing-value imputation, robust feature scaling, optional univariate feature selection or PCA, and multiple candidate classification or regression algorithms. Hyperparameters were optimized within a nested cross-validation framework with 10 inner and 10 outer folds (36). Balanced accuracy was used for sex classification, and R², mean squared error, and mean absolute error were used for regression targets. Full pipeline details and hyperparameter ranges are provided in Supplementary Methods S4.

To contextualize the information contained in hier-shPLRNN parameters, we included two baseline representations. First, we performed PCA on vectorized empirical fMRI time series and retained the same number of components as subject-specific hier-shPLRNN parameters. The resulting PCA scores served as a data-driven low-dimensional baseline (37). Second, we used empirical rs-fMRI FC directly as a high-dimensional reference representation by vectorizing the upper triangular elements of each empirical FC matrix. These baselines were entered into the same statistical and machine learning analyses as the hier-shPLRNN parameters.

## 3. Results

### 3.1 Hyperparameter Optimization

Hyperparameter optimization identified a hier-shPLRNN configuration with 20 subject-specific parameters and a shared hidden size of 1294 as optimal according to the validation criterion, defined as the mean Pearson correlation between empirical and simulated FC after subject-specific fine-tuning. This configuration was used for all subsequent analyses of reconstruction performance, generalization, subject specificity, and phenotypic associations. The full hyperparameter search is shown in Supplementary Figure S5.

### 3.2 Reconstruction performance across training, validation, and held-out test sets

Using the selected configuration, model performance was quantified on the training, validation, and held-out test sets using two complementary metrics: (i) Pearson correlation between vectorized empirical and simulated FC matrices and (ii) DTW distance between empirical and simulated multivariate time series.

On the training set (n = 945), the model achieved a mean FC correlation of r = 0.63 and a mean DTW distance of 108.8. On the validation set used for hyperparameter selection (n = 208), mean performance after subject-specific fine-tuning was r = 0.51 with a DTW distance of 112.7. On the held-out test set (n = 191), performance was comparable, with a mean FC correlation of r = 0.51 and a DTW distance of 112.5. These results are summarized in Figure 2. Examples of empirical and simulated time series and FC matrices for representative held-out test subjects are shown in Figure 3.

**Figure 2:**
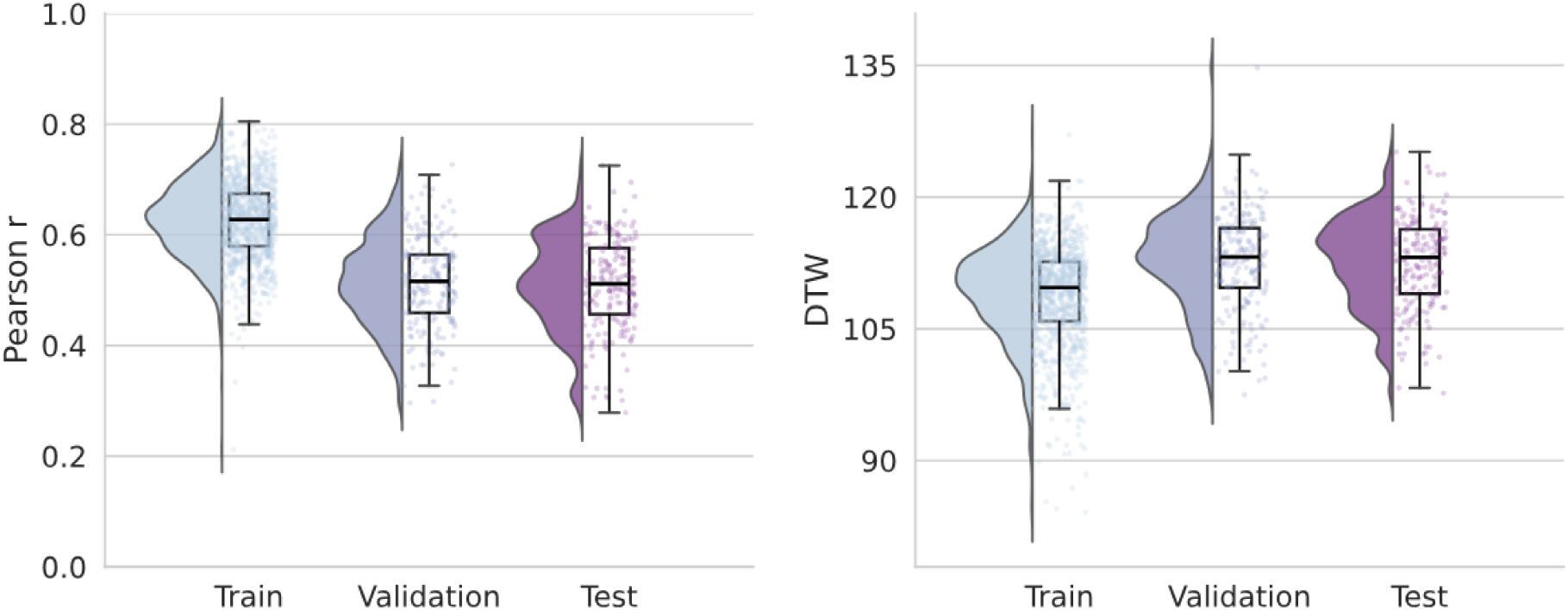
Reconstruction performance across datasets. Left: Pearson correlation between empirical and simulated FC matrices. Right: DTW distance between z-scored empirical and simulated time series. Training subjects were optimized jointly (shared + individual parameters). For validation and test subjects, only individual parameters were optimized during subject-specific fine-tuning while shared parameters were fixed.

**Figure 3:**
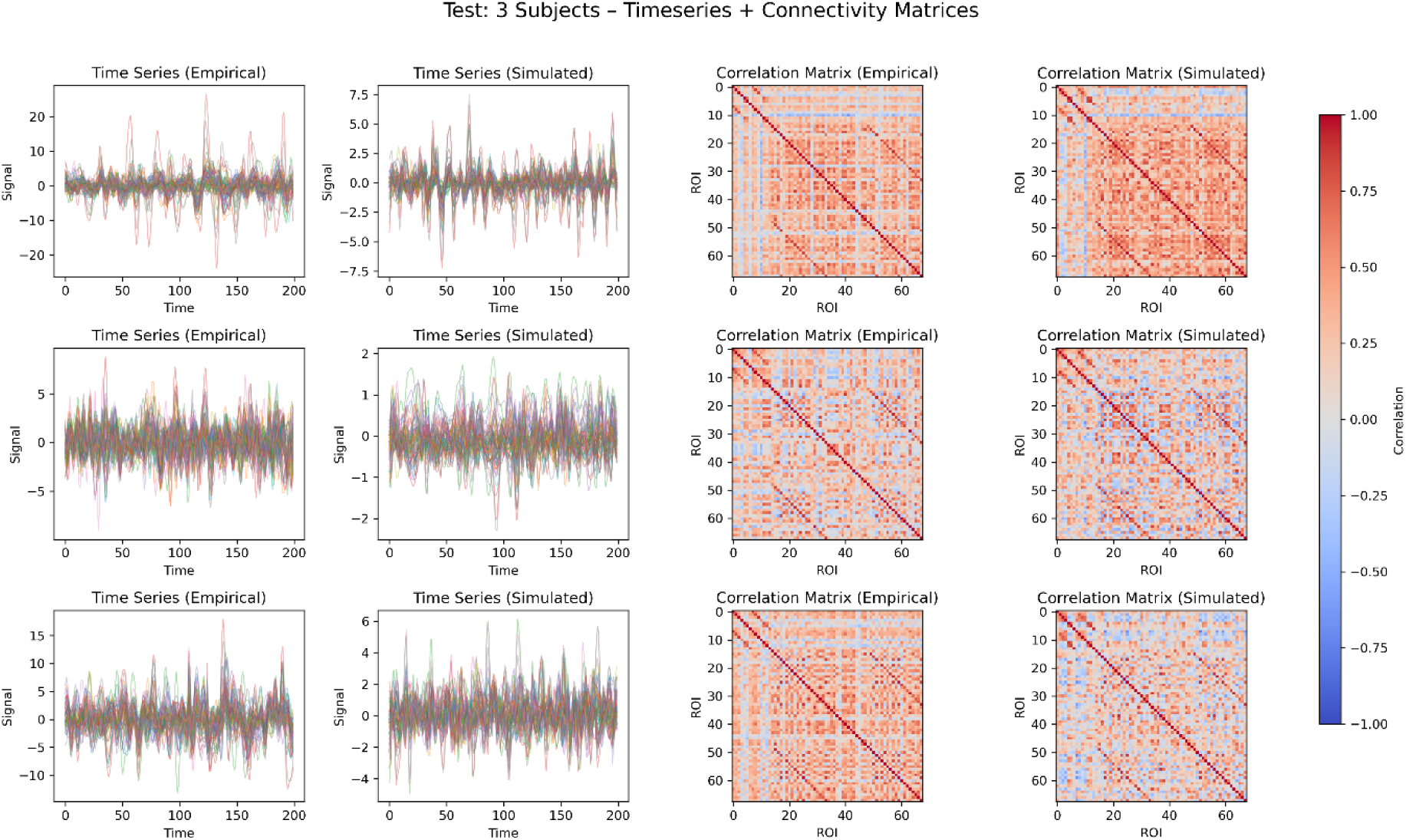
Simulated vs. empirical rs-fMRI time series for three exemplary test subjects after fine-tuning. Left: Time series of rs-fMRI activity across 68 brain regions and 200 timepoints, comparing empirical (left column) and simulated (right column) data for three test subjects. Right: Functional connectivity (FC) matrices based on pairwise Pearson correlations between 68 brain regions, shown for empirical (left column) and simulated (right column) rs-fMRI data. Each row corresponds to a different test subject.

### 3.3 Subject-wise variability and template similarity

To assess the contribution of population-level connectivity structure, we compared empirical and simulated FC patterns with the group-mean empirical FC of the training set. Across held-out test subjects (n = 191), empirical FC showed a mean correlation of r = 0.667 with the group template, exceeding the reconstruction accuracy of the individualized model (r = 0.510). Interestingly, simulated FC showed even greater similarity to the group template (r = 0.717), indicating that model-generated FC patterns were more strongly aligned with population-level connectivity structure than the empirical individual FC patterns themselves (Figure 4A).

**Figure 4.**
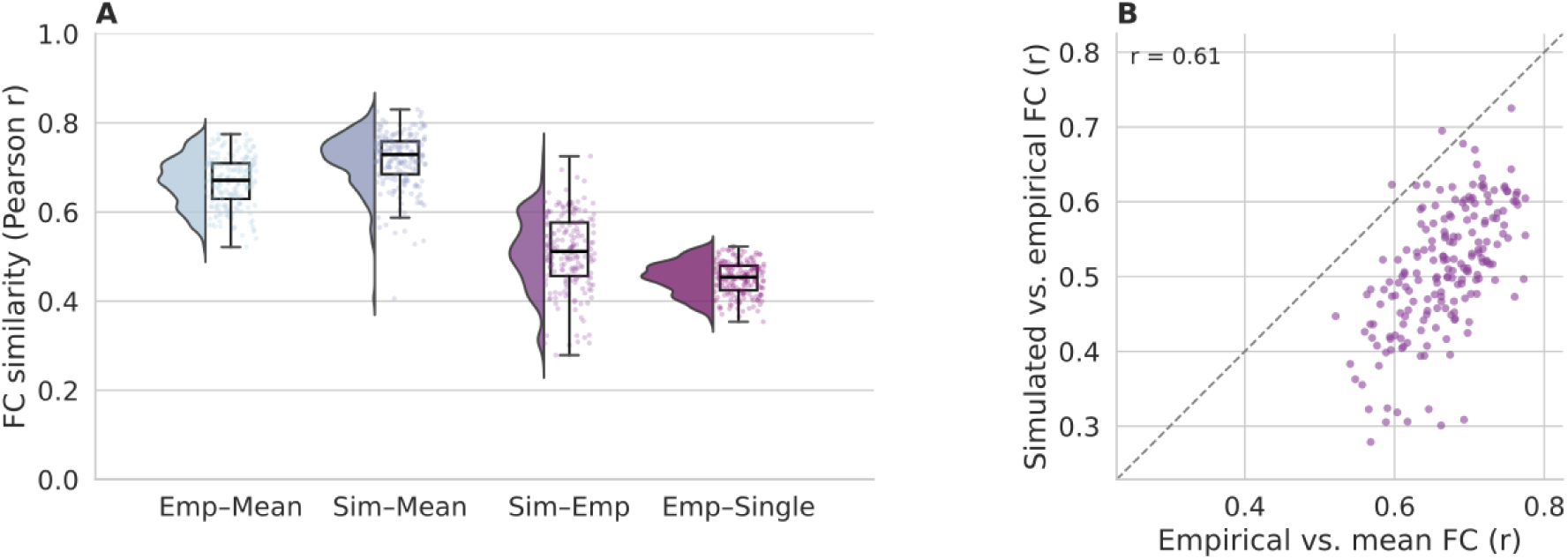
Population-level connectivity structure dominates individual FC reconstruction. (A) Distributions of Pearson correlations between individual empirical FC and the empirical group-mean FC (Emp–Mean), simulated FC and the empirical group-mean FC (Sim–Mean), simulated and empirical FC of the same subject (Sim–Emp), and empirical FC and the FC of single training subjects (Emp–Single; noise-matched baseline, averaged across all training subjects). (B) Relationship between empirical template similarity (Emp–Mean) and reconstruction accuracy (Sim–Emp) across held-out test subjects. Each point represents one subject. Empirical group-mean FC and single-subject FC were calculated exclusively from the training set (n = 945); all comparisons were performed on held-out test subjects (n = 191).

Note that the template averages FC across 945 scans and is therefore largely free of sampling noise, whereas each simulated FC derives from a single 200-time-point realization. A noise-matched baseline, defined as the mean correlation with the FC of each individual training subject, correlated with individual empirical FC at r = 0.451; the model exceeded this baseline in 83% of test subjects (Wilcoxon p < 10⁻²⁰). Simulated FC was also more similar to the subject’s own empirical FC than to that of other test subjects (r = 0.510 vs. 0.474, p < 10⁻⁹), although this subject-specific component was small.

Reconstruction accuracy varied substantially across held-out test subjects and increased with template similarity, defined as the correlation between a subject’s empirical FC and the group template (Figure 4B). A linear regression model with template similarity as the sole predictor explained R² = 0.37 of the variance in reconstruction accuracy. Thus, subjects whose empirical FC patterns were more similar to the training cohort template were reconstructed more accurately.

### 3.4 Subject identification based on FC

To assess whether simulated FC retained subject-specific information beyond group-level structure, we performed an FC-based subject identification analysis within the training set. For each subject, we correlated the simulated FC matrix with all empirical FC matrices and recorded the rank of the matching subject’s empirical FC.

The matching subject was ranked first in 96/945 cases (Top-1: 10.16%), among the three highest correlations in 178/945 cases (Top-3: 18.84%), and among the five highest correlations in 226/945 cases (Top-5: 23.92%). Chance-level Top-1 identification was approximately 0.1% (1/945). These results indicate that simulated FC retained subject-discriminative information above chance, although identification accuracy remained below what is typically expected from empirical FC fingerprinting (9). The full correlation matrix underlying the FC-based subject identification analysis is shown in Supplementary Figure S6.

### 3.5 Test-retest stability of subject-specific parameters

To assess test-retest stability, we analyzed 308 training subjects with two rs-fMRI sessions acquired approximately two years apart. Subject-specific parameter vectors were estimated independently for each session, while keeping group-level parameters fixed, and compared using Pearson correlation across the full 20-dimensional parameter vector.

Hier-shPLRNN parameters showed higher within-subject than between-subject similarity. Mean within-subject similarity was r = 0.286 ± 0.238, whereas mean between-subject similarity was r = 0.027 ± 0.240, yielding Δr = 0.259. This separation was significant (permutation test: p = 5 × 10⁻⁵; Mann-Whitney U test: p = 7.55 × 10⁻³³) and associated with large effect sizes (Cohen’s d = 1.083; Cliff’s δ = 0.553).

Empirical FC exhibited higher absolute test-retest similarity. Within-subject FC similarity was r = 0.613 ± 0.083, compared with between-subject similarity of r = 0.459 ± 0.078, yielding Δr = 0.154. This separation was also significant (permutation test: p = 5 × 10⁻⁵; Mann-Whitney U test: p = 4.63 × 10⁻⁷³) and associated with large effect sizes (Cohen’s d = 1.914; Cliff’s δ = 0.840). Thus, subject-specific model parameters were reproducible above between-subject similarity, but empirical FC remained the more stable representation in absolute terms (Figure 5).

**Figure 5.**
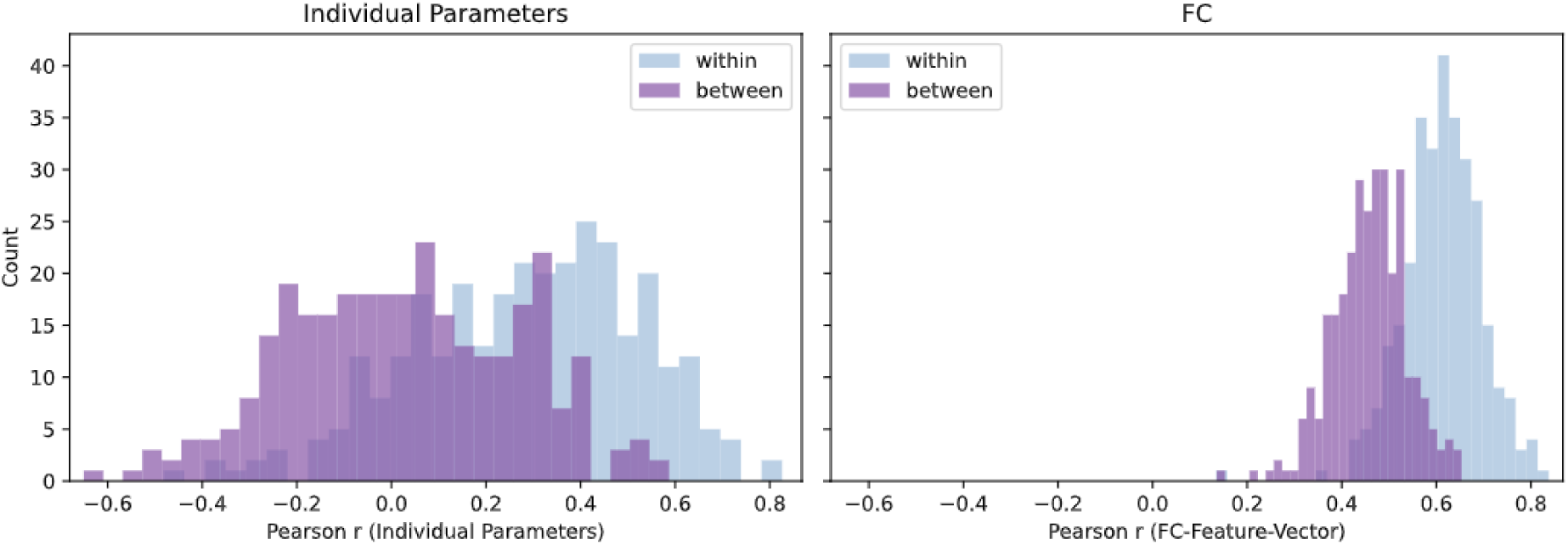
Test-retest similarity of individualized model parameters and functional connectivity. Left: Pearson correlation between subject-specific shPLRNN parameter vectors estimated from two rs-fMRI sessions, shown for within-subject and between-subject comparisons. Right: Corresponding test-retest similarity of empirical functional connectivity (FC), quantified as the Pearson correlation between vectorized FC matrices. Within-subject similarity exceeds between-subject similarity in both modalities, indicating subject-specific and reproducible representations.

Across subjects, parameter test-retest similarity was positively associated with FC test-retest similarity (Pearson r = 0.267, p = 2.02 × 10⁻⁶; Spearman r = 0.260, p = 3.89 × 10⁻⁶). This indicates that subjects with more stable empirical FC also tended to show more stable subject-specific model parameters.

### 3.6 Associations between Learned Parameters and Subject Characteristics

In univariate statistical analyses, hier-shPLRNN-derived parameters showed FDR-corrected associations with sex, age, and BMI, whereas associations with years of schooling and IQ did not survive correction. Effect sizes were modest, with a maximum partial 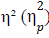 of 0.021 for sex and a maximum absolute standardized β (β*_std_*) of 0.372 for continuous outcomes. PCA-derived parameters yielded fewer significant associations and smaller effect sizes. Empirical rs-fMRI features showed stronger and more consistent associations across phenotypic variables (Table 1).

**Table 1:**
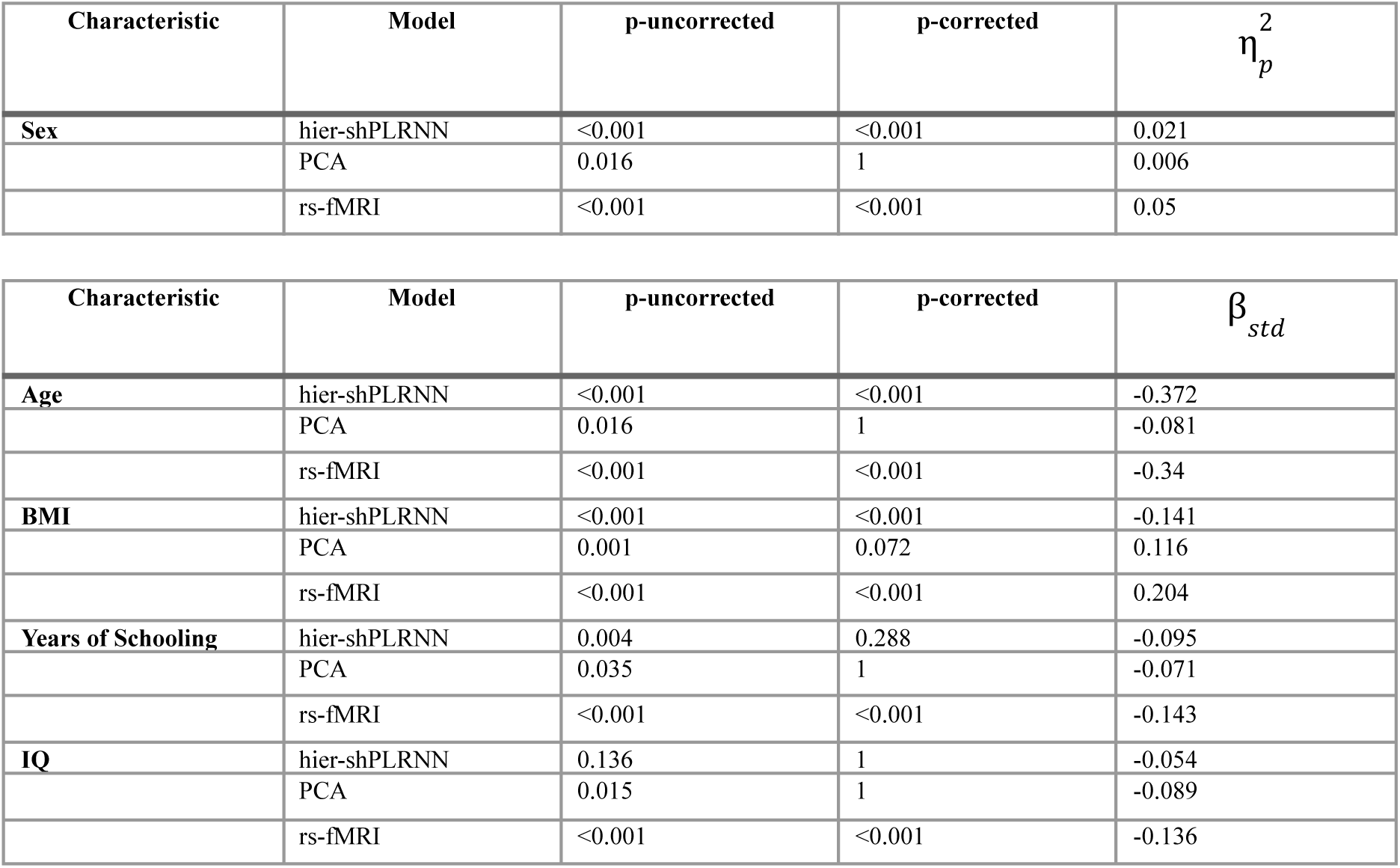
Univariate associations between feature representations and subject characteristics. Statistical analyses were performed separately for hier-shPLRNN-derived parameters, PCA-derived parameters, and empirical rs-fMRI features. Sex was analyzed using ANOVA, while age, BMI, years of schooling, and IQ were analyzed using OLS regression. Reported values correspond to the strongest feature-level association for each target and representation after correction for multiple comparisons.

| **Characteristic** | **Model** | **p-uncorrected** | **p-corrected** | η<sub>p</sub><sup>2</sup> |
|---|---|---|---|---|
| **Sex** | hier-shPLRNN | <0.001 | <0.001 | 0.021 |
|  | PCA | 0.016 | 1 | 0.006 |
|  | rs-fMRI | <0.001 | <0.001 | 0.05 |
| **Characteristic** | **Model** | **p-uncorrected** | **p-corrected** | β<sub>std</sub> |
| **Age** | hier-shPLRNN | <0.001 | <0.001 | -0.372 |
|  | PCA | 0.016 | 1 | -0.081 |
|  | rs-fMRI | <0.001 | <0.001 | -0.34 |
| **BMI** | hier-shPLRNN | <0.001 | <0.001 | -0.141 |
|  | PCA | 0.001 | 0.072 | 0.116 |
|  | rs-fMRI | <0.001 | <0.001 | 0.204 |
| **Years of Schooling** | hier-shPLRNN | 0.004 | 0.288 | -0.095 |
|  | PCA | 0.035 | 1 | -0.071 |
|  | rs-fMRI | <0.001 | <0.001 | -0.143 |
| **IQ** | hier-shPLRNN | 0.136 | 1 | -0.054 |
|  | PCA | 0.015 | 1 | -0.089 |
|  | rs-fMRI | <0.001 | <0.001 | -0.136 |

### 3.7 Machine learning prediction of subject characteristics

We next assessed whether subject-specific parameters contained multivariate predictive information about subject characteristics. Across all machine learning models evaluated with hier-shPLRNN-derived parameters as input, the best sex classification model achieved a balanced accuracy of BACC = 0.70. For regression targets, the best-performing models achieved MSE values of 76.89 for age, 12.40 for BMI, 6.04 for years of schooling, and 142.52 for IQ (Table 2).

**Table 2:** Machine learning prediction of subject characteristics from different feature representations. Performance of the best-performing machine learning pipeline for each target variable using hier-shPLRNN-derived parameters, PCA-derived parameters, or empirical rs-fMRI features as input. Sex classification was evaluated using balanced accuracy (BACC). Regression targets were evaluated using mean squared error (MSE), mean absolute error (MAE), and explained variance.

| **Characteristic** | **Model** | **BACC** |  |  |
|---|---|---|---|---|
| **Sex** | hier-shPLRNN | 0.7 |  |  |
|  | PCA | 0.64 |  |  |
|  | rs-fMRI | 0.78 |  |  |
| **Characteristic** | **Model** | **MSE** | **MAE** | **explained variance** |
| **age** | hier-shPLRNN | 76.89 | 6.76 | 0.49 |
|  | PCA | 137.09 | 10.17 | 0.01 |
|  | rs-fMRI | 59.41 | 6.23 | 0.6 |
| **BMI** | hier-shPLRNN | 12.4 | 2.82 | 0.18 |
|  | PCA | 15.09 | 3.12 | 0.006 |
|  | rs-fMRI | 9.04 | 2.4 | 0.39 |
| **Years of Schooling** | hier-shPLRNN | 6.04 | 2.16 | 0.05 |
|  | PCA | 6.34 | 2.15 | 0.005 |
|  | rs-fMRI | 6.03 | 2.16 | 0.05 |
| **IQ** | hier-shPLRNN | 142.52 | 10.03 | 0.04 |
|  | PCA | 151.2 | 10.48 | -0.01 |
|  | rs-fMRI | 145.06 | 10.37 | 0.05 |

Hier-shPLRNN-derived parameters consistently outperformed PCA-derived low-dimensional representations. Empirical rs-fMRI features provided the strongest performance for sex, age, and BMI prediction, while performance for years of schooling and IQ was comparable across representations and generally weak. Overall, these results suggest that hier-shPLRNN parameters retain multivariate information about individual characteristics, but do not outperform empirical rs-fMRI features as predictive representations.

## 4. Discussion

In this study, we systematically evaluated a hierarchical shPLRNN framework for individualized dynamical system reconstruction from resting-state fMRI. The model combines shared group-level recurrent dynamics with compact subject-specific parameters, aiming to generate individual rs-fMRI trajectories whose long-term properties reproduce empirical FC. Our results show that hierarchical recurrent dynamical models can recover substantial FC structure from short rs-fMRI time series and generalize to held-out subjects after subject-specific fine-tuning. At the same time, the analyses reveal clear limitations: reconstruction accuracy depended strongly on the similarity of a subject’s FC profile to the group template, empirical FC remained the stronger reference for static subject identification and prediction, and associations between learned parameters and phenotypic variables were modest.

A first methodological finding concerns the scaling behavior of the model. Hyperparameter optimization selected a large shared hidden layer, consistent with scaling principles previously proposed for hierarchical dynamical system reconstruction in lower-dimensional benchmark systems (19,20). This suggests that modeling high-dimensional empirical neuroimaging data requires substantial shared recurrent capacity, and that the size of this shared backbone can be guided by principled scaling considerations rather than purely ad hoc tuning. In contrast, increasing the number of subject-specific parameters did not improve performance. The optimal model used only 20 subject-specific parameters, and larger individual parameter spaces did not yield better reconstruction accuracy. This indicates that, in the present setting, individual differences were best captured by a compact representation embedded within a sufficiently expressive shared dynamical model. More subject-specific degrees of freedom therefore did not automatically translate into better personalization and may instead increase sensitivity to noise or weaken the regularizing effect of the shared group-level structure.

Using the optimized configuration, the model achieved higher reconstruction accuracy in the training set than in unseen subjects, but performance after subject-specific fine-tuning was comparable between the validation and held-out test sets, suggesting that the model generalized consistently once shared group-level dynamics had been learned. However, reconstruction accuracy varied substantially across individuals, and a large portion of this variability was explained by template similarity: subjects whose empirical FC more closely resembled the group-mean FC of the training cohort were reconstructed more accurately, whereas atypical subjects were more difficult to model. The template-based comparisons further clarify this dependence. The empirical group-mean FC approximated individual empirical FC more closely than the model reconstructions did, and simulated FC was even more similar to the group template than empirical FC itself. Together, this indicates that the model learned a dominant population-level dynamical structure and captured individual deviations from it only to a limited extent. Nevertheless, simulated FC exceeded a noise-matched single-subject baseline and retained a small but significant subject-specific component beyond the template.

These findings highlight an important distinction between reproducing shared connectivity structure and recovering individual-specific dynamics. While the group-mean FC provides a strong static baseline, it does not constitute a generative dynamical model. From a dynamical systems perspective, the goal of the hier-shPLRNN is not to reproduce empirical BOLD time series point by point, but to learn a generative process whose autonomous trajectories preserve relevant long-term properties of the observed system, here primarily FC. The model’s ability to generate trajectories with realistic FC supports its usefulness as a generative representation of rs-fMRI. Strengthening individualization, and robustness to atypical or out-of-distribution connectivity patterns, will be an important direction for future work. Longer or repeated recordings and averaging FC over multiple simulated realizations may help in this respect.

We further assessed whether the learned subject-specific parameters captured stable individual information. Test-retest analyses showed that parameter vectors were more similar within subjects than between subjects across sessions acquired approximately two years apart. This indicates that the learned low-dimensional representations retain reproducible subject-specific information. However, empirical FC showed higher absolute test-retest similarity and stronger within-between separation. Thus, hier-shPLRNN parameters should not be interpreted as replacing empirical FC as a static subject fingerprint. Rather, they provide a compact dynamical representation that retains some subject-specific structure while being constrained by a shared generative model. The positive association between FC test-retest stability and parameter stability further suggests that the reliability of learned parameters partly depends on the reliability of the underlying empirical FC structure.

Subject identification analyses led to a similar conclusion. Simulated FC enabled above-chance identification of subjects from the training set, indicating that generated trajectories retained subject-discriminative information. However, identification accuracy remained limited compared with what is typically observed for empirical FC fingerprinting. This comparison highlights an important distinction between empirical FC and generative dynamical modeling. Empirical FC is a direct, high-dimensional summary of covariance structure and is therefore expected to be a strong static fingerprint. In contrast, the hier-shPLRNN compresses individual information into a low-dimensional parameter vector embedded in a shared dynamical system.

Associations between subject-specific parameters and demographic or psychometric variables were statistically detectable but modest. In univariate analyses, hier-shPLRNN parameters were associated with sex, age, and BMI, but effect sizes were small, and empirical rs-fMRI features generally showed stronger associations. Multivariate machine learning analyses confirmed that hier-shPLRNN parameters contained predictive information, particularly for sex and age, and outperformed PCA-based low-dimensional representations. However, empirical rs-fMRI features remained the strongest representation for most targets. These findings are consistent with prior work showing that rs-fMRI contains phenotypic information, but that effect sizes are often modest and prediction performance depends strongly on the representation used (7). They also suggest that the learned dynamical parameters capture only part of the individual variability present in empirical FC.

Several limitations should be considered. First, the study was based on a single cohort and one cortical parcellation. We used the 68-region Desikan–Killiany atlas to obtain stable, low-dimensional ROI time series for dynamical modeling, but this choice may limit sensitivity to fine-grained individual functional topography. The hierarchical shPLRNN framework is not inherently restricted to anatomical parcellations and could, in principle, also be applied to functionally derived atlases, such as Schaefer or Glasser, as well as finer-grained parcellations. However, increasing the number of ROIs also increases the dimensionality of the modeled system and the associated computational demands. With 200 time points per sample, a 400-region parcellation would yield more observed dimensions than time points, so finer parcellations may require longer recordings, stronger regularization, or a latent dimensionality below the number of observed regions (i.e., a non-identity observation model). Our training-size analysis (Supplement S2) further suggests that several hundred subjects are needed for transferable group-level dynamics even at 68 regions. Moreover, finer parcellations may yield noisier regional signals, potentially increasing the data requirements for reliable estimation of individual dynamics. While such extensions are technically feasible in principle, their practical performance, particularly in smaller datasets, remains to be established. Future work should therefore systematically investigate the trade-off between parcellation granularity, model complexity, and subject-specific reconstruction performance. Second, all time series were truncated to 200 time points. This enabled standardized model input across subjects but may limit the complexity of individual dynamics that can be reliably inferred. Longer recordings may provide stronger constraints on subject-specific dynamical parameters. Third, censored frames were removed and the remaining frames concatenated rather than interpolated. This introduces occasional discontinuities that violate the temporal continuity assumed by dynamical systems reconstruction. We expect the impact to be limited for several reasons: (i) Censoring was conservative: frames were removed only when both FD and DVARS exceeded their thresholds. Splice points therefore affect at most 1.6% of one-step transitions; (ii) Censoring was applied after band-pass filtering, so splice points join already smoothed signal values; (iii) The primary evaluation metric, FC, is insensitive to frame order; (iv) Reconstruction accuracy was not meaningfully associated with the number of censored frames, and uncensored samples did not outperform censored ones (Supplement S7). The alternatives carry their own drawbacks. Interpolation inserts synthetic data that the model would learn as genuine dynamics. Requiring contiguous motion-free segments would substantially reduce the sample and bias it towards low-motion participants. Longer or multi-echo acquisitions could avoid this trade-off in future work. Fourth, the model was evaluated against PCA-based representations and empirical FC, but not against a broader set of deep generative time-series models. Direct comparisons with alternative recurrent, variational, or generative sequence models would be useful to clarify which findings are specific to the hierarchical shPLRNN architecture. Fifth, the model showed reduced performance for subjects with atypical FC profiles, suggesting limited robustness to out-of-distribution dynamics. Finally, although associations with phenotypic variables were statistically significant for some targets, their effect sizes were modest, limiting immediate interpretability in behavioral or clinical terms.

In summary, hierarchical shPLRNNs provide a principled framework for learning compact generative representations of individual rs-fMRI dynamics. The model recovered substantial FC structure, generalized to held-out subjects after subject-specific fine-tuning, and produced low-dimensional parameters that were reproducible across sessions and informative about some subject characteristics. At the same time, empirical FC remained the stronger static representation for subject identification and phenotypic prediction, and model performance depended strongly on the similarity of individual FC profiles to the group template. These findings support hierarchical dynamical modeling as a promising approach for individualized rs-fMRI, while also clarifying its current limits: the model captures individual dynamics primarily as deviations around a shared population-level structure, and further work is needed to improve sensitivity to atypical and fine-grained individual differences.

## Acknowledgments

This work was funded in part by the consortia grants from the German Research Foundation (DFG) FOR 2107, SFB/TRR 393 (“Trajectories of Affective Disorders”, project grant no 521379614), and the Germany’s Excellence Strategy (EXC 3066/1 “The Adaptive Mind”, Project No. 533717223), as well as the DYNAMIC center, funded by the LOEWE program of the Hessian Ministry of Science and Arts (grant number: LOEWE1/16/519/03/09.001(0009)/98).

During the preparation of this work the authors used ChatGPT to improve language and readability. After using this tool, the authors reviewed and edited the content as needed and take full responsibility for the content of the published article.

## Data Availability Statement

The MACS data is not publicly available due to participant consent and data protection restrictions. Data may be made available from the corresponding author upon reasonable request and subject to institutional and ethical approval.

## Code Availability Statement

Analysis code is available from the corresponding author upon reasonable request.

