## Supplements for "Promise and Limits of Hierarchical Dynamical RNNs for Individualized Resting-State fMRI"

#### S1: Influence of individual parameters

**Methods.** To see how much influence the individual parameters have in our setting, we ran a hier-shPLRNN with only one individual parameter per subject. All other parameters remain the same as before. Performance was quantified using (i) the Pearson correlation between each subject's empirical and simulated functional connectivity (FC) matrices, and (ii) the dynamic time warping (DTW) distance between empirical and simulated multivariate time series.

**Results.** The mean Pearson correlation between simulated and empirical rs-fMRI was r = 0.29. The mean dynamic time warping (DTW) was 118.3 (vs. r = 0.63 and DTW = 108.8).

**Discussion.** Reducing the number of subject-specific parameters from 20 to 1 markedly reduced both FC reconstruction accuracy and temporal similarity. This indicates that subject-specific parameters are essential for capturing individual FC structure and individual temporal dynamics.

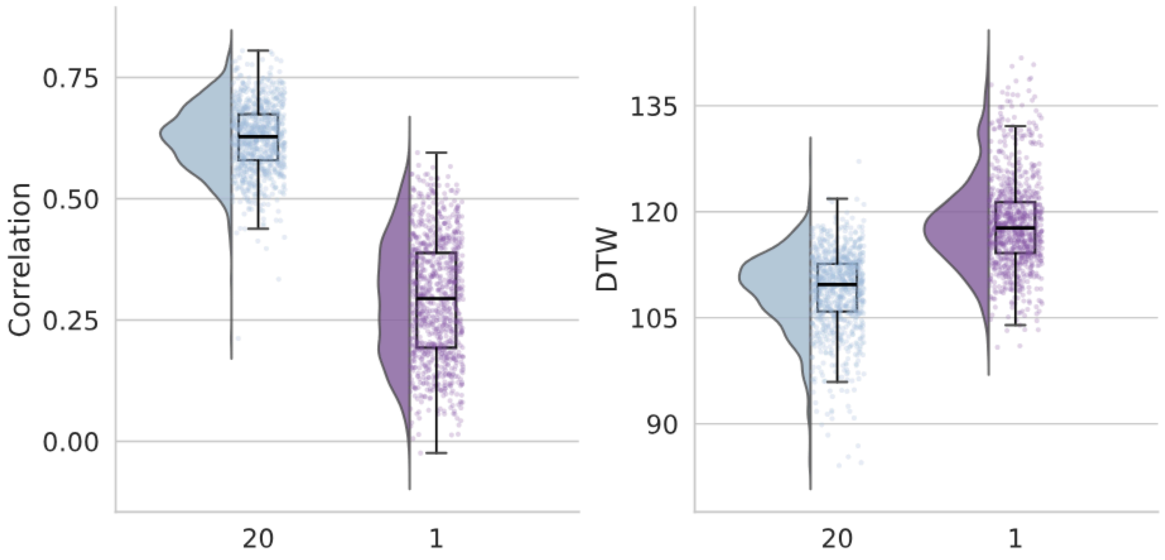


**Figure S1: Pearson Correlation & Dynamic Time Warping for 20 and 1 individual parameter.** Values for Pearson correlation between simulated and empirical FC (left) and DTW distance between z-scored empirical and simulated time series (right). For '20' the train set (945 subjects) was optimized with 20 individual parameters; for '1' with only one individual parameter.

#### S2: Effect of training cohort size on reconstruction accuracy

**Methods.** To assess how training set size affects model performance and generalization, we re-trained the hier-shPLRNN on subsets of the data with different numbers of subjects (50, 100 and 500) and compared the results to the model trained on the full training cohort (n = 945). All models used the same architecture and optimization procedure. After training, performance was evaluated within each condition by comparing simulated and empirical data for the respective subjects. Performance was quantified using (i) the Pearson correlation between each subject's empirical and simulated functional connectivity (FC) matrices, and (ii) the dynamic time warping (DTW) distance between empirical and simulated multivariate time series.

To assess generalization, we additionally performed subject-specific fine-tuning on the held-out test set for each model variant. During fine-tuning, only the subject-specific parameters were updated, while the group-level dynamics learned from the respective training subset (e.g., 50, 100 or 500 subjects) were kept fixed. Reconstruction performance on the test set was then quantified using the same FC correlation and DTW metrics.

**Results.** Training set size influenced reconstruction performance in two partially dissociable ways.

(i) Fit to the training cohort. When each model was evaluated on the same cohort it was trained on, all models achieved moderate to high correspondence between empirical and simulated FC. Mean Pearson correlations ranged from $r = 0.40$ to $r = 0.69$ across training sizes (50 subjects: $r = 0.40$; 100: $r = 0.41$; 500: $r = 0.69$; 945: $r = 0.63$). DTW distance did not show a systematic dependence on training set size (50: 110.3; 100: 111.9; 500: 107.3; 945: 108.8).

(ii) Generalization to new subjects. To evaluate generalization, group-level dynamics were frozen and subject-specific fine-tuning was performed on the held-out test set. After this adaptation step, reconstruction accuracy on unseen subjects depended more strongly on the size of the training cohort. Models trained on 50 and 100 subjects achieved relatively low FC correspondence after fine-tuning ($r = 0.29$ and $r = 0.32$, respectively), whereas models trained on 500 subjects reached substantially higher performance ($r = 0.52$). The model trained on the full cohort (945 subjects) achieved comparable FC reconstruction accuracy ($r = 0.51$). DTW values on the test set were similar across training sizes (111.5-115.5).

See Supplementary Figure S2 and S3 for details.

**Discussion.** Taken together, these results indicate that training on small cohorts was sufficient to fit FC structure within the same subjects but yielded less transferable group-level dynamics. Training on several hundred subjects substantially improves generalization to unseen individuals after subject-specific fine-tuning, particularly in terms of FC reconstruction, while gains beyond approximately 500 subjects appear to level off. Temporal alignment as measured by DTW was largely independent of training set size and therefore less discriminative for generalization performance than FC.

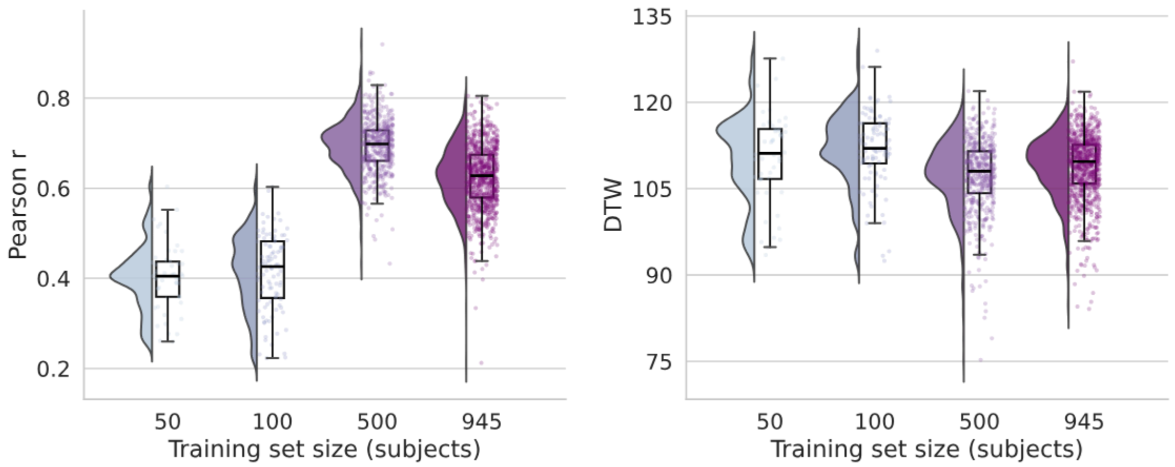


**Supplementary Figure S2. Effect of training cohort size on reconstruction accuracy.** Pearson correlation between empirical and simulated FC (left) and DTW distance between z-scored empirical and simulated time series (right) for models trained on different numbers of subjects (50, 100, 500, 945), evaluated on the corresponding training cohort.

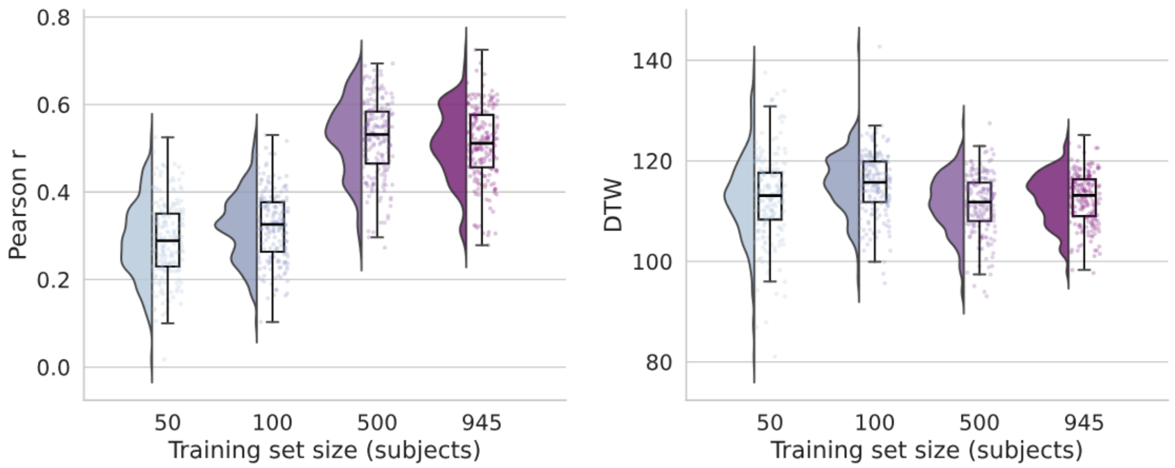


**Supplementary Figure S3. Effect of training cohort size on reconstruction accuracy and generalization.** Pearson correlation between empirical and simulated FC (left) and DTW distance between z-scored empirical and simulated time series (right) for models trained on different numbers of subjects (50, 100, 500, 945), finetuned and evaluated on the held-out test cohort.

### S3: Run-to-run reliability of subject-wise functional connectivity

**Methods.** We quantified how reproducibly the model generates subject-specific FC across independent training runs. We trained the hier-shPLRNN five times from scratch under the same settings. For each run and each subject, we simulated time series from the trained model and computed the subject's FC matrix (Pearson correlation across time between all region pairs). We then vectorized the upper triangle of each FC matrix (excluding the diagonal) and, for every subject, computed the Pearson correlation between the FC estimates obtained from two different runs. This yielded a subject-wise FC correlation for each run pair; we then averaged these correlations across subjects.

**Results.** Run-to-run FC correlations were consistently high. Mean subject-wise FC correlation across runs ranged from r = 0.73 to r = 0.79 for all run pairs (Supplementary Figure S4). This indicates that the subject-specific FC patterns generated by the model are highly reproducible across independent optimizations of the model.

**Discussion.** The high between-run reproducibility suggests that the hier-shPLRNN converges to stable subject-level FC structure: even when the model is re-trained from scratch, the simulated FC for a given subject remains highly similar. This stability supports that the individual differences the model learns are not arbitrary artifacts of a particular training run but reflect consistent features the model can recover across runs.

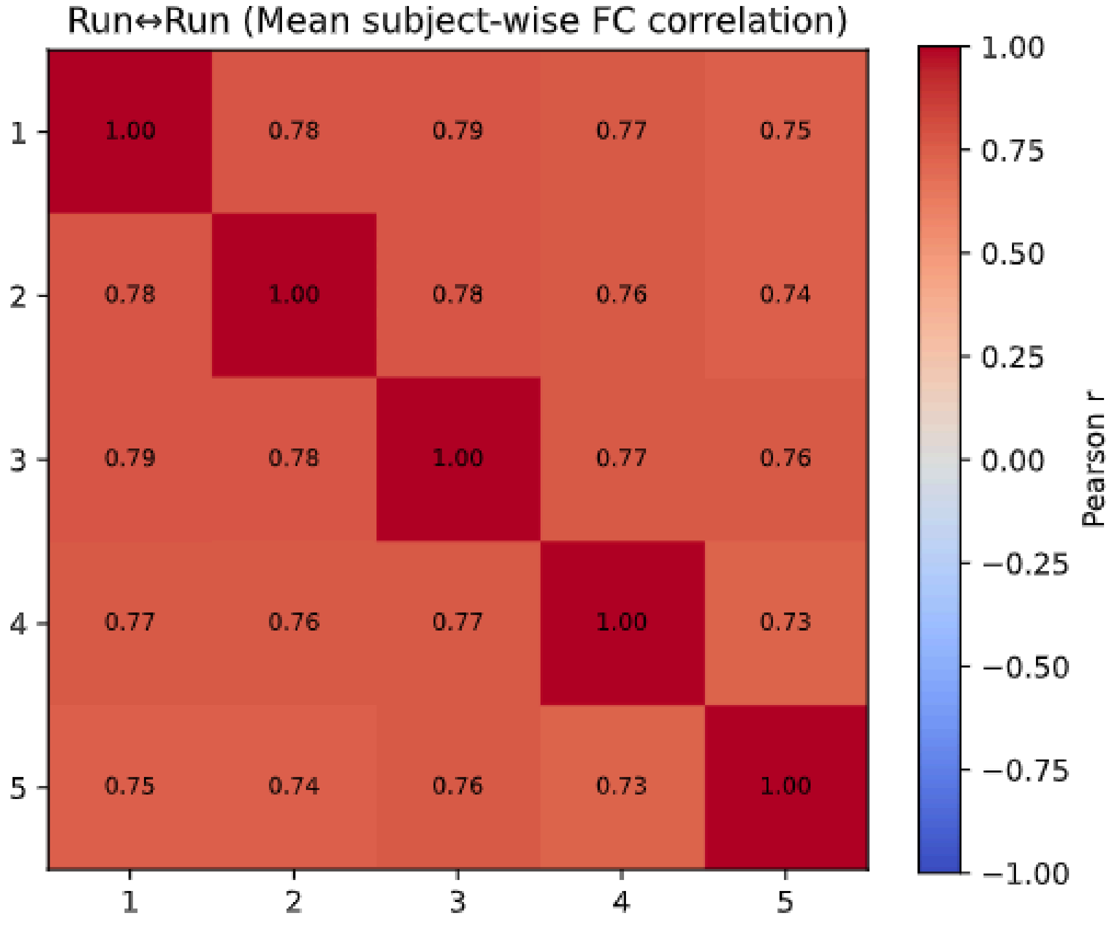


**Supplementary Figure S4. Run-to-run reproducibility of subject-specific functional connectivity (FC) estimates.** The matrix shows the mean Pearson correlation of subject-wise FC between pairs of independent model training runs (Runs 1-5). For each subject and each run, we computed the FC matrix from the model-generated time series after training, vectorized its upper triangle, and correlated it with the corresponding FC from another run for the same subject. Values shown are averaged across all subjects. Off-diagonal entries quantify between-run reproducibility; diagonal entries are 1.0 by definition. All off-diagonal mean correlations were high (r = 0.73-0.79), indicating that the model yields stable subject-level FC structure across independent training runs.

**S4: Machine Learning Analysis**

To assess whether subject-specific model parameters contained multivariate information about individual characteristics, we performed supervised machine learning analyses using PHOTONAI. Analyses were conducted for sex, age, body mass index (BMI), years of schooling (YoS), and intelligence quotient (IQ). To avoid data leakage across repeated measurements, only the first available session per subject was included. Subjects with missing values for the respective target variable were excluded from the corresponding analysis.

Machine learning analyses were performed separately for three input representations: (i) subject-specific feature vectors estimated by the hierarchical shPLRNN, (ii) PCA-based low-dimensional representations of empirical fMRI time series, and (iii) empirical rs-fMRI functional connectivity (FC). For the PCA baseline, each subject's fMRI time series was vectorized across time points and ROIs, and the number of retained principal components was matched to the number of subject-specific parameters in the hierarchical shPLRNN. For the empirical FC baseline, FC matrices were vectorized by extracting the upper triangular elements excluding the diagonal.

All analyses used nested cross-validation with 10 outer folds and 10 inner folds. The outer cross-validation loop was used to estimate generalization performance, while the inner loop was used for hyperparameter optimization and model selection. All preprocessing steps were fitted only on the respective training folds and then applied to the corresponding validation or test folds to prevent data leakage.

Each machine learning pipeline consisted of feature scaling, optional dimensionality reduction or feature selection, and a supervised learning algorithm. A mean-imputation step was included as a safeguard but did not affect the analyses because the final input matrices contained no missing feature values. Features were scaled using a RobustScaler to reduce the influence of outliers. Dimensionality reduction and feature selection were treated as alternative preprocessing options using a PHOTONAI Switch. Candidate options included univariate feature selection based on ANOVA F-values, retaining 5%, 10%, or 50% of features, and principal component analysis (PCA) using full variance decomposition.

For sex classification, candidate algorithms included support vector machines, random forests, logistic regression, k-nearest neighbors, Gaussian naïve Bayes, and gradient boosting classifiers. Logistic regression models were evaluated with L1, L2, and elastic-net penalties. Support vector machines were evaluated with linear, polynomial, and radial basis function kernels across a range of regularization strengths. For continuous targets, corresponding regression algorithms were evaluated, including support vector regression, random forest regression, k-nearest-neighbor regression, and gradient boosting regression. Hyperparameters were optimized within the inner cross-validation loop.

Classification performance was quantified using balanced accuracy to account for potential class imbalance. Regression performance was quantified using explained variance, mean squared error (MSE), and mean absolute error (MAE). Reported performance values correspond to the mean performance across the outer cross-validation folds. The same nested cross-validation structure, preprocessing options, model families, and performance metrics were used for all input representations to ensure comparability between hierarchical shPLRNN parameters, PCA-based representations, and empirical FC.

### S5: Hyperparameter Optimization

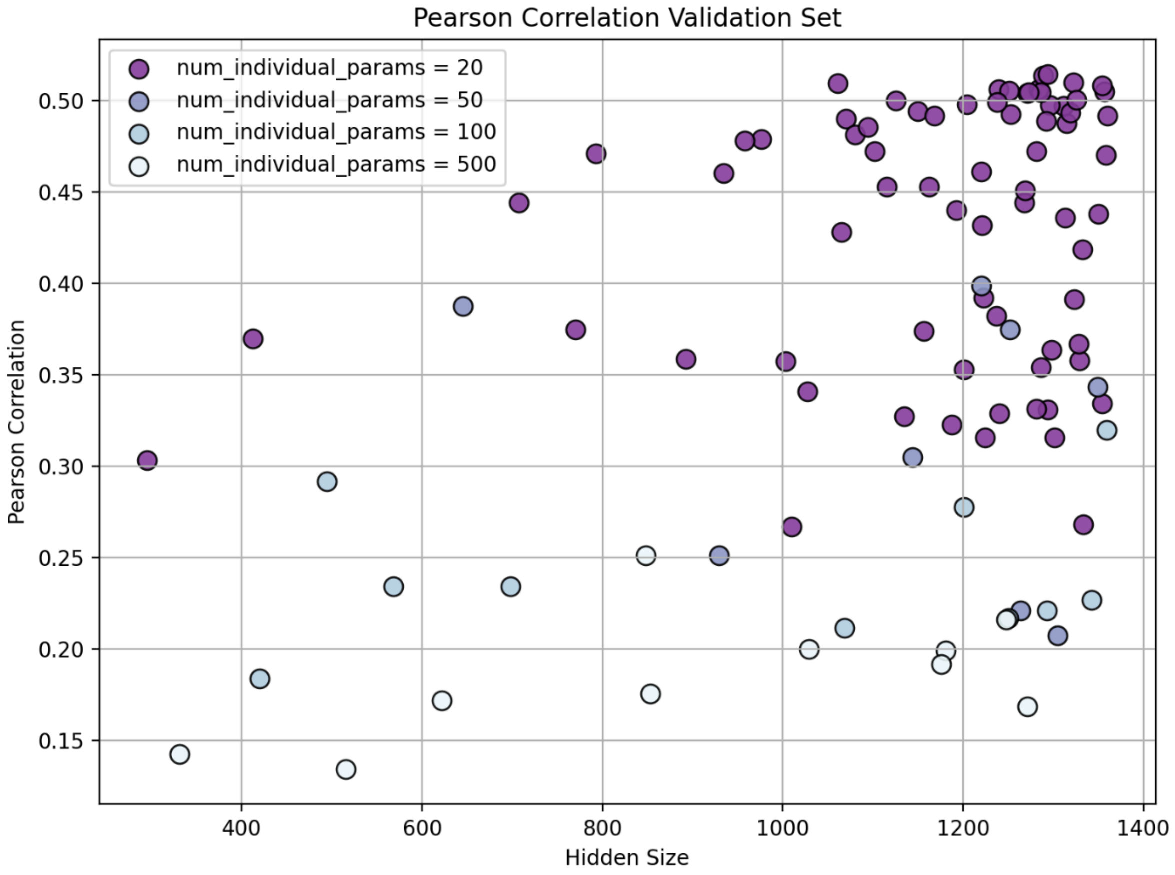


**Supplementary Figure S5: Result of hyperparameter optimization for hidden size and number of individual parameters.** Hyperparameter optimization results across hidden size and number of individual parameters. Each point represents one Optuna trial (training on 945 subjects and subsequent fine-tuning on 208 validation subjects; 1000 epochs each). Color indicates the number of individual parameters, and the y-axis shows the Pearson correlation between empirical and simulated FC after fine-tuning. The best configuration (20 individual parameters, hidden size = 1294) achieved r = 0.51.

### S6: FC-based subject identification

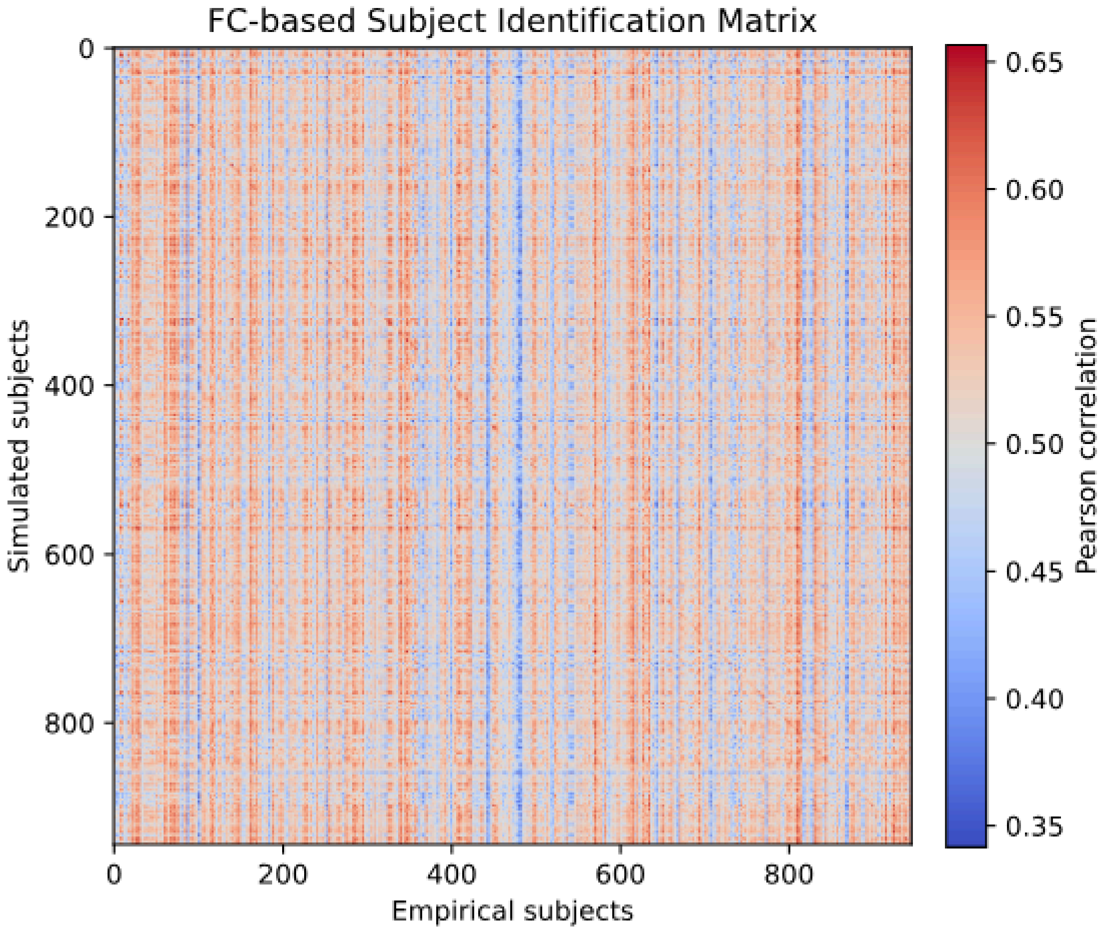


**Supplementary Figure S6. FC-based subject identification matrix.** Matrix of Pearson correlations between simulated functional connectivity (FC) matrices (rows) and empirical FC matrices (columns) for all subjects in the training set (n = 945). Diagonal elements correspond to same-subject comparisons. While elevated correlations are observed along the diagonal, substantial off-diagonal structure indicates limited subject discriminability, consistent with the reported Top-1, Top-3, and Top-5 identification accuracies.

### S7: Influence of motion censoring on reconstruction accuracy

**Methods.** Frames exceeding motion thresholds were removed and the remaining frames concatenated (Section 2.2). The resulting splice points violate the temporal continuity assumed by dynamical systems reconstruction. To assess whether this affected model performance, we determined the number of removed frames for each sample (0-23 of 223 frames). We related it to per-subject FC reconstruction accuracy and DTW distance using Spearman correlation. In addition, we compared samples without any censored frames to samples with at least one censored frame (Mann-Whitney U test; bootstrap 95% confidence intervals of the mean difference, 10,000 resamples; Cliff's δ). Because censoring was associated with higher empirical signal variance, we repeated the analyses while adjusting for each sample's mean empirical signal standard deviation (log-transformed; linear regression). All analyses used the final model configuration and were performed separately for the training, validation, and test sets.

**Results.** Across all samples, 23.3% contained no censored frames. The median number of removed frames was 5 (95th percentile: 19; maximum: 23). FC reconstruction accuracy was not associated with the number of removed frames in any data split, and uncensored samples did not show higher accuracy than censored samples (Table S1). After adjusting for signal variance, a small negative association emerged in the training set (−0.0009 per removed frame, $p = 0.007$; i.e., −0.02 at the maximum of 23 frames) but not in the validation or test sets (both $p > 0.07$). DTW distances were lower (indicating higher similarity) in censored than in uncensored samples (Table S1). In the validation and test sets, this difference was fully explained by signal variance (adjusted $p > 0.6$). In the training set, a small association remained (−0.11 per removed frame, $p < 0.001$).

**Discussion.** We found no evidence that the discontinuities introduced by frame removal impaired reconstruction accuracy. Associations were either absent or of negligible magnitude, and differences in DTW pointed in the opposite direction to what discontinuities would predict. This is consistent with the small fraction of affected one-step transitions (at most 1.6%; Section 2.2).

| | **Train** | **Validation** | **Test** |
|---|---|---|---|
| n (uncensored) | 945 (222) | 208 (49) | 191 (42) |
| **FC r** | | | |
| ρ (removed frames, r_FC) | 0.01 (p=.71) | -0.08 (p=.28) | 0.05 (p=.52) |
| Uncensored/censored | 0.619/0.627 | 0.518/0.514 | 0.490/0.516 |
| 95% CI of difference | [-0.018, 0.002] | [-0.018, 0.027] | [-0.056, 0.003] |
| Cliff’s δ | -0.07 | 0.03 | -0.15 |
| Adjusted for signal SD | -0.0009 (p=.007) | -0.0015 (p=.08) | -0.0008 (p=.43) |
| **DTW** | | | |
| ρ (removed frames, DTW) | -0.22 (p<.001) | -0.04 (p=.55) | -0.16 (p=.02) |
| Uncensored/censored | 110.4/108.3 | 114.2/112.2 | 114.8/111.9 |
| 95% CI of difference | [1.3, 2.8] | [0.5, 3.4] | [1.5, 4.4] |
| Cliff’s δ | 0.22 | 0.19 | 0.33 |

| | | | |
|---|---|---|---|
| Change per removed frame, adjusted for signal SD | -0.11 ($p<.001$) | 0.02 ($p=.68$) | 0.00 ($p=.99$) |

**Supplementary Table S1. Association between motion censoring and reconstruction accuracy.** Results are shown separately for the training, validation, and test sets using the final model configuration. n (uncensored): number of samples in total and number of samples without any censored frames. ρ: Spearman correlation between the number of removed frames (0–23) and per-sample FC correlation ($r_{FC}$) or DTW distance. Uncensored / censored: mean accuracy of samples without vs. with at least one censored frame. 95% CI of difference: bootstrap confidence interval (10,000 resamples) of the mean difference (uncensored − censored). Cliff's δ: non-parametric effect size of the same group comparison; positive values indicate higher values in uncensored samples. Change per removed frame, adjusted for signal SD: regression coefficient for the number of removed frames when controlling for the log-transformed mean empirical signal standard deviation. FC accuracy is the Pearson correlation between simulated and empirical FC (higher = better). DTW is the distance between z-scored empirical and simulated time series, per time point (lower = better).
